# Syllable-PBWT for space-efficient haplotype long-match query

**DOI:** 10.1101/2022.01.31.478234

**Authors:** Victor Wang, Ardalan Naseri, Shaojie Zhang, Degui Zhi

## Abstract

The positional Burrows-Wheeler transform (PBWT) has led to tremendous strides in haplotype matching on biobank-scale data. For genetic genealogical search, PBWT-based methods have optimized the asymptotic runtime of finding long matches between a query haplotype and a predefined panel of haplotypes. However, to enable fast query searches, the full-sized panel and PBWT data structures must be kept in memory, preventing existing algorithms from scaling up to modern biobank panels consisting of millions of haplotypes. In this work, we propose a space-efficient variation of PBWT named Syllable-PBWT, which divides every haplotype into syllables, builds the PBWT positional prefix arrays on the compressed syllabic panel, and leverages the polynomial rolling hash function for positional substring comparison. With the Syllable-PBWT data structures, we then present a long match query algorithm named Syllable-Query. Compared to Algorithm 3 of Sanaullah et al. (2021), the most time- and space-efficient previously published solution to the long match query problem, Syllable-Query reduced the memory use by a factor of over 100 on both the UK Biobank genotype data and the 1000 Genomes Project sequence data. Surprisingly, the smaller size of our syllabic data structures allows for more efficient iteration and CPU cache usage, granting Syllable-Query even faster runtime than existing solutions. The implementation of our algorithm is available at https://github.com/ZhiGroup/Syllable-PBWT.

## 1 Introduction

Developments in genotyping technologies have accelerated the growth of genotype databases, paving the way for systematically comparing the haplotype sequences inherited by individuals [1, 2]. Shared, long DNA segments, known as Identical by Descent (IBD) segments, between the haplotypes of two or more individuals are highly indicative of a recent common ancestor [3]. To efficiently analyze large panels of haplotypes, Durbin proposed the positional Burrows-Wheeler transform (PBWT) [4], a data structure that reorders haplotypes at every site (i.e. position within a haplotype) to concisely represent local substring matches within sets of aligned haplotypes, and has a construction runtime proportional to the size of the panel. Beyond IBD segment detection [5, 6, 7], PBWT has found applications in genotype imputation [8, 9], studying recombination events [10], and inferring ancestral recombination graphs [11].

PBWT algorithms come in two flavors: finding all within-panel pairwise haplotype matches (all-vs-all matching), or finding all pairwise haplotype matches between an out-of-panel haplotype and any in-panel haplotype (one-vs-all query). In this work, we are concerned with the query problem, of which an important application is genealogical search. Durbin’s Algorithm 5 [4] is able to find all set-maximal matches from a query haplotype to any panel haplotype, where a set-maximal match is said to exist from a haplotype *s*_1_ to a haplotype *s*_2_ if no other haplotype in the panel has a longer match with *s*_1_ that completely contains the range of sites over which *s*_1_ and *s*_2_ match. However, as noted in [12], reporting only set-maximal matches is likely to exclude a lot of valuable match information, since many considerably long matches would not be reported simply because they were overshadowed by a longer match. (Note too that the quality of being set-maximal is not necessarily symmetric; i.e. that a match is set-maximal from *s*_1_ to *s*_2_ does not imply that it is set-maximal from *s*_2_ to *s*_1_, which is unintuitive for genealogical search.) Instead, setting a match length cutoff is more theoretically justifiable and has been the common practice in real-world genealogical search deployed by direct-to-consumer (DTC) genetics companies. In spite of the errors present in haplotype data, DTC genetics companies and other researchers have demonstrated the efficacy of using long matches to determine genealogical relationships [13, 14, 12]. In the PBWT-Query work [12], Naseri et al. defined an *L*-long match (abbreviated to “long match”) to be a match spanning at least *L* sites (or, for genetic distance, at least *L* cM) and presented an algorithm to find all long matches between a query haplotype and a panel in average-case *O*(*N* + *c*) time, where there are *N* sites and *c* reported matches. Remarkably, since *O*(*N*) time is indispensable to read in the query haplotype, and *O*(*c*) to output matches, *O*(*N* + *c*) is the fastest time complexity theoretically achievable.

However, existing PBWT query algorithms are not space-efficient. Although all-vs-all PBWT matching consumes minimal memory as the scanning algorithms only store data relevant to the current site, one-vs-all PBWT query entails retaining data for all sites in memory to enable efficient pointer lookups when iterating through the sites. To bypass previously visited matches and achieve efficient runtime, Naseri et al. introduced data structures called LEAP arrays, which increase the memory burden, on top of what is already required by the original PBWT data structures. To lighten memory usage, Sanaullah et al. developed Algorithms 3 and 4 of d-PBWT [15], which solve the long match query problem without LEAP arrays in worst-case and average-case runtimes, respectively, of *O*(*N* + *c*). Despite the memory improvement, storing PBWT data structures in memory for the whole genome remains a bottleneck for potential applications, such as online wholegenome query services. For example, to query on the 22 autosomal chromosomes from UK Biobank consisting of 974,818 haplotypes and 658,720 markers, Algorithms 3 and 4 of Sanaullah et al. require 10.1 TB of memory. Accommodating memory usage of this magnitude demands dedicating expensive servers with massive amounts of RAM. Moreover, even the size of UK Biobank’s database pales in comparison to the tens of millions of genotype samples collected by DTC companies, and this number is only set to rise [16].

For servers with relatively limited memory, current alternatives include keeping data on the SSD or HDD, often in tandem with memory-mapped files. However, accessing these sources is accompanied by a significant runtime overhead, which, when memory-mapped files are used, also heavily depends on the similarity between previous and subsequent queries, as discussed by the authors of PBWT-Query [12]. Alternatively, distributing the PBWT panel into multiple servers may lower the memory footprint for individual servers but at the incurred cost of synchronization.

In this work, we present a space-efficient variation of the PBWT data structure, named Syllable-PBWT, with which we in turn present Syllable-Query, an algorithm that solves the *L*-long match query problem with more optimal memory usage and runtime than existing algorithms. One theoretical contribution featured in this work is the replacement of the divergence array, which in past works has gone hand in hand with the PBWT data structure, with polynomial hashing. While the basic idea of chunking into syllables is core to our approach, the innovation mainly lies in our adaptation of PBWT algorithms, which traditionally were geared towards bi-allelic (or at best multi-allelic) sequences, to function on general sequences.

## 2 Methods

### 2.1 Overview

The existing algorithms for the *L*-long match query problem, as presented by Naseri et al. [12] and Sanaullah et al. [15], use the binary haplotype sequences to construct the PBWT, which we refer to as bit-PBWT. To query with bit-PBWT, said algorithms maintain three full-panel-sized (comprising *MN* integers) data structures: the positional prefix arrays *a*, the divergence arrays *d*, and the virtual extension arrays *u* and *v*. We reasoned that the dense encoding by bit-PBWT would be redundant for identifying *L*-long matches where *L* is large, since short matches can simply be skipped. Thus, we propose Syllable-PBWT, which treats every *B* contiguous sites as one syllable and builds data structures for only every syllable rather than for every site. In doing so, Syllable-PBWT reduces the size of positional prefix arrays by a factor of *B*. Further, Syllable-PBWT introduces prefix hash arrays to eliminate and replace the divergence arrays and virtual insertion arrays. To further reduce the panel size, we perform coordinate compression and build dictionaries at each syllable, leveraging the linkage disequilibrium of the haplotype sequences. Overall, the space usage of Syllable-PBWT is about *B* times smaller than that of bit-PBWT, as outlined in Table 1. In order to identify all *L*-long matches, we develop the Syllable-Query algorithm using the Syllable-PBWT data structure. The following subsections elaborate upon the presented algorithms and their correctness, and the pseudocode for construction and query is given in Appendix 6.1 and 6.2, respectively.

**Table 1:** Space comparison between bit-PBWT and Syllable-PBWT data structures used to query in Algorithm 3 of Sanaullah et al. and Syllable-Query, respectively. Values are in units of *MN* 32-bit integers, where there are *M* haplotypes with *N* sites each (assume *N* is a multiple of B). *ρ* ≥ 1 is defined in Section 2.3.1. The hashes are stored as 64-bit integers, hence the 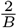 memory from *h*.

|  |  | Bit-PBWT | Syllable-PBWT |
| --- | --- | --- | --- |
| Panel of sequences | $\dot{X}/X$ | $\frac{1}{32}$ | $\frac{1}{B}$ |
| Syllable dictionaries | $r$ | - | $\frac{1}{32\rho}$ |
| Positional prefix arrays | $a$ | 1 | $\frac{1}{B}$ |
| Prefix hash arrays | $h$ | - | $\frac{2}{B}$ |
| Divergence arrays | $d$ | 1 | - |
| Virtual extension arrays | $u, v$ | 2 | - |
| Total space | | $4 + \frac{1}{32}$ | $\frac{4}{B} + \frac{1}{32\rho}$ |

### 2.2 Notation

The data we are dealing with is a haplotype panel consisting of aligned binary haplotype sequences. In a sequence *s*, we denote the value at position *b* as *s*[*b*] (the first value is *s*[0]), and *s*[*b*, *e*) denotes the sequence of values from position *b* to position *e* – 1, inclusive. An *L*-long match (abbreviated to “match”) between sequences *s*_1_ and *s*_2_ is said to start at *b* and end at *e* if *s*_1_[*b*, *e*) = *s*_2_[*b*, *e*), *s*_1_[*b* – 1] ≠ *s*_2_[*b* – 1] (or *b* = 0), *s*_1_[*e*] ≠ *s*_2_[*e*] (or *e* is the length of the sequences), and *e* – *b* ≥ *L* for some specified *L*. Let 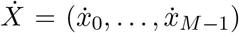 be the panel of *M* haplotype sequences, each with *N* sites, with which queries are to be matched. Off of the haplotype panel 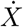, we will construct a raw syllabic panel 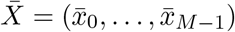 and, in turn, a (compressed) syllabic panel *X* = (*x*_0_,…, *x*_*M* – 1_). The construction and the length *n* of every raw/compressed syllabic sequence is later described. For any collection of sequences *C* = (*c*_0_,…, *c*_*M*–1_), we let *C*[*k*] = (*c*_0_[*k*],…, *c*_*M*–1_[*k*]).

### 2.3 Syllable-PBWT

The Syllable-PBWT data structure consists of the syllabic panel *X* with dictionaries *r*, the positional prefix arrays *a*, and the polynomial prefix hash arrays *h*.

#### 2.3.1 Syllabic panel

To shorten the length of the sequences, we split the panel into syllables of *B* sites each, padding the ends of the haplotypes with 0s so that the number of sites becomes a multiple of *B*. For the *k*th *B*-site syllable of the ith haplotype, we parse the binary allele values spanning the *B* sites, i.e. the reverse of *ẋ_i_*[*kB*, (*k* + 1)*B*), as a binary number, whose value we assign to 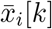, syllable *k* of the raw syllabic sequence 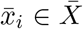. Constructing 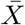 takes *O*(*MN*) time since it is computationally equivalent to reading in the panel.

Although we have reduced the length of the sequences by a factor of *B* to get 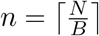 syllables, our raw syllabic panel 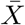 still contains the same underlying information, merely arranged into *B*-bit integers, as 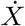. To reduce the space required to store our syllabic sequences, we observe that the number of distinct raw syllable values at a given syllable is bounded by the number of haplotypes *M*. If i*M* << 2^*B*^, we can apply coordinate compression (i.e. mapping sparse values in a large space to dense values in a small space) to the raw syllable values at a given syllable to obtain the compressed syllable values (abbreviated to “syllable values”). To enable conversion between raw and compressed syllable values, we build *r_k_*, a sorted dictionary of the distinct raw syllable values at syllable *k*. Then, every (compressed) syllabic sequence *x_i_* ∈ *X* can be built as follows: *x_i_*[*k*] is the index of 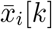 in *r_k_*, where said index can be found with binary search. The second step of Figure 1 illustrates the compression. The raw syllable values can later be recovered using the dictionary: 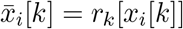.

**Figure 1:**
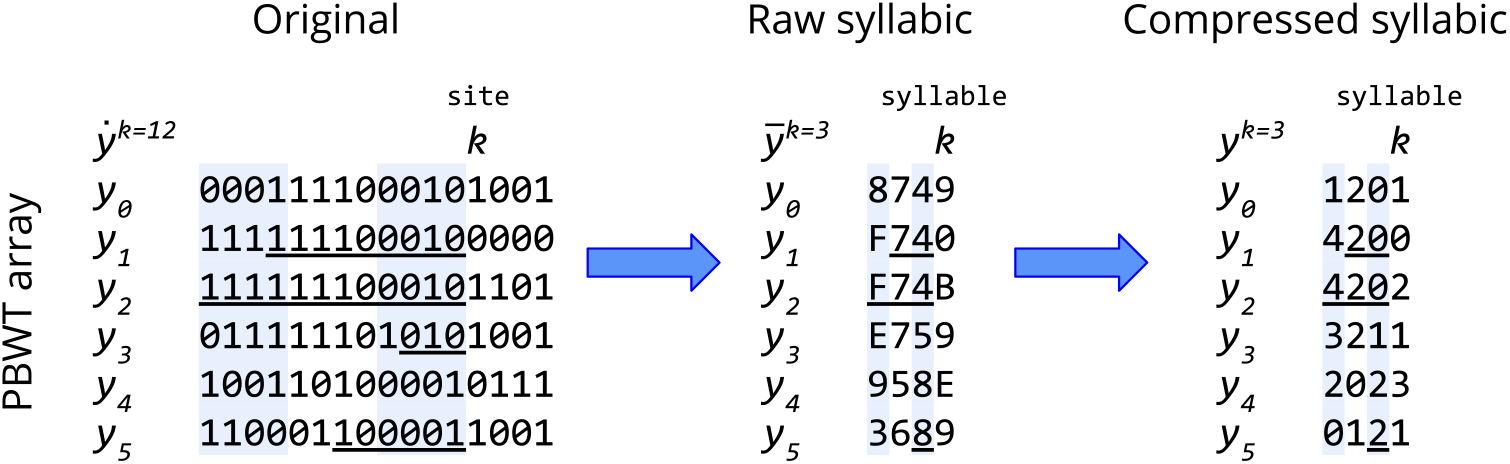
The reverse of every *B* = 4 binary allele values parsed as a binary number to obtain raw syllable values, written in hexadecimal (*A* = 10,…, *F* = 15), which undergo coordinate compression to produce the compressed syllable values. Underlines indicate the reverse prefix match before site/syllable *k* with the preceding sequence in the positional prefix order *a_k_*.

Since 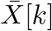 can be written in terms of *r_k_* and *X*[*k*], we can avoid the redundancy of keeping 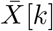 in memory after precomputation on syllable *k*. Instead, we store *X* with *O*(*Mn*) memory and *r* with *O*(*B*|*r*|) memory. In the worst case, in which at every syllable, all the sequences have distinct syllable values, then |*r*| = *Mn* implies *r* will require *O*(*MnB*) = *O*(*MN*) memory. Fortunately, in genetic data, linkage disequilibrium (i.e. non-random association of alleles across sites [17]) gives rise to repetitive syllable values at any given syllable. Therefore, the ratio 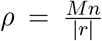 will likely be considerable, and *r* will use 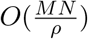 memory. Due to the sorting and binary search on the raw syllable values to compute *r* and *X*, respectively, they each take *O*(*Mnβ*log *M*) time to compute, for some small factor *β* ∈ *O*(*B*); due to the efficiency of 64-bit architectures, *β* << *B* (see Appendix 6.3 for details).

#### 2.3.2 Positional prefix array and PBWT array

The positional prefix array *a_k_* serves as the backbone of PBWT by storing the ordering of the sequences’ reverse prefixes before position *k*. In other words, for the syllabic panel *X*, the position of *i* in *a_k_* is the rank of the reverse of *x_i_*[0, *k*) when sorted (in lexicographical order) among, for all *j*, the reverse of *x_j_*[0, *k*). To simplify notation, the PBWT array *y^k^* is defined such that 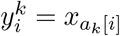, i.e. the sequence at position *i* in *a_k_* (*ẏ* and *ȳ* are similarly defined according to 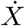 and 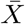, respectively); the *y* arrays need not be kept in memory as they can be expressed in terms of *X* and *a*. Algorithm 1 of Durbin’s PBWT makes use of the binary nature of allele values in bit-PBWT so that two pointers can be used to build *a*_*k*+1_ in *O*(*M*) time, given *a_k_* and 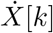.

To build *a*_*k*+1_ for sequences with up to *M* possible syllable values in Syllable-PBWT, we employ similar reasoning to that in bit-PBWT. Figure 1 visualizes the syllabic PBWT array in relation to the binary PBWT array. *X*[*k*] is the most significant syllable in determining the ordering of *a*_*k*+1_. If two sequences have the same value at syllable *k*, then the tie is to be broken with their reverse prefixes over syllables [0, *k*). In other words, *a*_*k*+1_ can be calculated by sorting the sequences by their *X*[*k*] value and, for ties, retaining the ordering from *a_k_*. This can be accomplished in *O*(*M*) time and memory with counting sort, a stable sorting algorithm, since the syllable values are bounded by *M*. Therefore, the positional prefix arrays over the *n* syllables require *O*(*Mn*) time and memory to compute and store.

#### 2.3.3 Polynomial prefix hash function and array for substring matching

The polynomial rolling hash function [18] is a simple and efficient hash function for substring matching. However, its applications in bioinformatics seem limited [19, 20]. One of our main observations is that the divergence arrays are not the only efficient bookkeeping method for positional substring matching in PBWT. The polynomial rolling hash function too can efficiently check if a pair of aligned sequences match over an interval. Specifically, the polynomial rolling hash function of the first *k* elements of *x_i_* is defined as

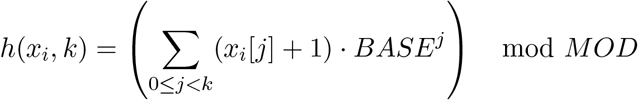

where *BASE* and *MOD* are distinct large primes [21, 18]. In other words, the expression within parentheses is obtained by adding 1 to every syllable value and parsing the reverse of the resulting *x_i_*[0, *k*) as a number in base *BASE*. The benefit of this hash function is that the hash value for any positional substring *x_i_*[*j*, *k*) can be calculated by *h*(*x_i_*, [*j*, *k*)) = *h*(*x_i_*, *k*) – *h*(*x_i_*, *j*) mod *MOD*. Thus, the polynomial rolling hash enables efficient substring matching given the prefix hashes. For justification on the dependability of our hash function despite possible collisions (less than 10^−9^ probability of collision over 10^10^ lookups), see Appendix 6.4. For notation, we define the polynomial prefix hash array *h_i_* such that *h_i_*[*k*] = *h*(*x_i_*, *k*) and *h_i_*[*j*, *k*) = *h*(*x_i_*, [*j*, *k*)).

Every syllable value is added to exactly one prefix hash exactly once, since every hash can simply build off of the previous syllable’s hash. Therefore, the arrays *h* require *O*(*Mn*) time and memory to compute and store.

### 2.4 Syllable-Query

Using the Syllable-PBWT data structures described above (*X*, *r*, *a*, *h*), we present the Syllable-Query algorithm to find long matches between a query haplotype and the panel. Crucial to Syllable-Query will be the hash arrays *h*, designed to replace the data structures *d*, *u*, *v* used for virtual insertion and finding matches. For the binary query haplotype sequence *ż*, we let 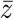 be its raw syllabic sequence and *z* be its (compressed) syllabic sequence. In the case that 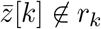, we let *z*[*k*] be a value distinct from the other *X*[*k*] values, such as the size of *r_k_*. We define *h_M_* by the same hash function as above according to *z*. Similarly to before, after reading in the query haplotype in *O*(*N*) time, these sequences require *O*(*nβ* log *M*) time to compute.

#### 2.4.1 Virtual insertion of query haplotype into panel without *u* and *v*

To enable out-of-panel query, past solutions have introduced the idea of virtually inserting the query haplotype into the panel [12]. The virtual locations of the query sequence are stored in an array *t* such that *t_k_* is the position in *a_k_* in which the query sequence would be, had it been included in the original panel. To calculate *t_k_*, past solutions utilize the precomputed auxiliary arrays *u* and *v* at every site *k* to facilitate computing *t*_*k*+1_ based on *ż*[*k*] and *t_k_*, where *u_k_*[*i*] is the number of 0 ≤ *j* < *i* for which 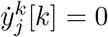, and *v_k_*[*i*] is *u_k_*[*M*] plus the number of 0 ≤ *j* < *i* for which 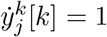. Specifically, *t*_*k*+1_ is *u_k_*[*t_k_*] if *ż*[*k*] =0 and *v_k_*[*t_k_*] otherwise. However, past solutions require binary sequences, and the notions of *u* and *v* do not efficiently generalize to *M* possible syllable values.

To find the value of *t*_*k*+1_, we binary search among the in-panel sequences for where *z* belongs. To compare *z* with another sequence *x_i_* in one step of the sequence-wise binary search, we first compare their values at syllable *k*, and if they are equal, we binary search for the maximum *b* < *k* for which *z*[*b*] ≠ *x_i_*[*b*] to compare *z* and *x_i_*. Once again, these fast comparisons are enabled by our hash arrays *h*. Our worst-case time complexity for virtual insertion over the *n* syllables is *O*(*n* log *M* log *n*), since we binary search over *M* sequences and *n* syllables. In the average case, the O(log *n*) time binary search would only occur in the small proportion of comparisons for which *z*[*k*] = *x_i_*[*k*] (the expected number of such comparisons is *ρ*), and the range of the syllable-wise binary search can be minimized by setting its lower bound to the greatest value of *b* when *t_k_* was being computed, since it is impossible for the start of the longest reverse prefix match with *z* at syllable *k* + 1 to be earlier than that at syllable *k*.

#### 2.4.2 Identifying long matches virtually near *z*

We define *l* as the minimum number of full syllables within any *L*-site match. To derive the expression for *l*, we must consider the case in which a match extends far into a syllable without completely covering it, i.e. *B* – 1 out of the *B* sites. If the remaining *L* – (*B* – 1) sites are to minimize the number of full syllables covered, they would not complete the nearly filled syllable but rather extend in the opposite direction. The number of full syllables covered would then be 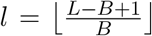. We consider matches spanning *l* syllables to be potential long matches, which we will abbreviate to “long matches” or “matches” with the implication that only matches spanning *L* sites after refinement will be reported; using bitwise operations on the raw syllable values, we can refine the single-site resolution boundaries for the 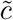 potential long matches in 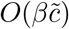 time (see Appendix 6.5 for details).

In Syllable-Query, we search for ongoing long matches, as opposed to past solutions’ focus on terminated matches, for reasons explained in Appendix 6.6. The definition of the positional prefix array guarantees that the sequences with the longest ongoing matches with a sequence 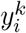 at syllable *k* occur in a contiguous block around position *i* in *a_k_*. Thus, at syllable *k*, we can iterate upwards and downwards within *a_k_* starting from *t_k_* until no more long matches are available. Since the process for finding matches above *z* is analogous to that below *z*, we will only describe the process for finding matches above *z* with the implication that a similar process is performed for matches below *z* (“above” and “below” refer to relative positions in the positional prefix array, with position 0 at the top and position *M* at the bottom).

To search for matches above *z*, we maintain a pointer *p* in *a_k_*. When there are no ongoing matches above *z*, we set *p* = *t_k_* – 1, and every time a match above *z* is found, *p* is decremented. We check for a match between *z* and 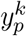 by checking whether *h_M_*[*k* – *l*, *k*) = *h*_*a*_*k*[*p*]__[*k* – *l*, *k*). Once this is false, there are no more matches above *z* to be found as of the current syllable *k*. Alternatively, we can cut back on the number of hash comparisons by binary searching for the final value of *p* (i.e. the block of new matches) before linearly iterating through the matches.

#### 2.4.3 Avoiding redundant counting of matches

From the process for identifying matches described above, it is evident that a match spanning *s* > *l* syllables will be counted *s* – *l* + 1 times, since that is the number of syllables *k* for which the matching sequence and *z* will match over syllables [*k* – *l*, *k*). If a query yields matches with an average length significantly greater than the minimum length *l*, then the runtime would suffer. Thus, we seek to count every match exactly once.

##### Lemma 1.

*Once a match with sequence x_1_ is identified immediately above z, sequence x_i_ must remain immediately above z until the match ends*.

Using Lemma 1 (see Appendix 6.7 for the proof), we can avoid redundantly visiting a match immediately above *z* at every syllable *k* < *m* ≤ *k* – *l* + *s*, after identifying it for the first time at syllable *k*, by preemptively setting our pointer *p* to *t_m_* – 2 rather than reconsidering the match. We further observe that Lemma 1 and its accompanying optimization can be generalized to any number of ongoing matches above *z*. That is, we maintain a running counter *up_on_*, and at every syllable *k*, we set *p* = *t_k_* – 1 – *up_on_* and every time another match is identified above *z*, we decrement *p* and increment *up_on_*, thereby bypassing previously identified matches.

The remaining task is to decrement *up_on_* every time we reach the end of a previously ongoing match. Let us maintain an array *up_end_* such that *up_end_*[*k*] stores the number of ongoing matches that end at syllable *k*, so that we can reduce *up_on_* by *up_end_*[*k*] before looking for matches at syllable *k*. To keep *up_end_* updated, we must find the total match length s of every match we identify and increment *up_end_*[*k* – *l* + *s*]. To do so efficiently, we binary search for the end of the match, checking whether *h_M_*[*k*, *m*) = *h*_*a*_*k*_[*p*]_[*k*, *m*) to test if a syllable *m* is a valid match end. Figure 2 depicts the process for finding long matches. Since there are *n* syllables over which we potentially must binary search, the runtime of extending the 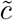 potential matches is 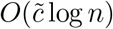.

**Figure 2:**
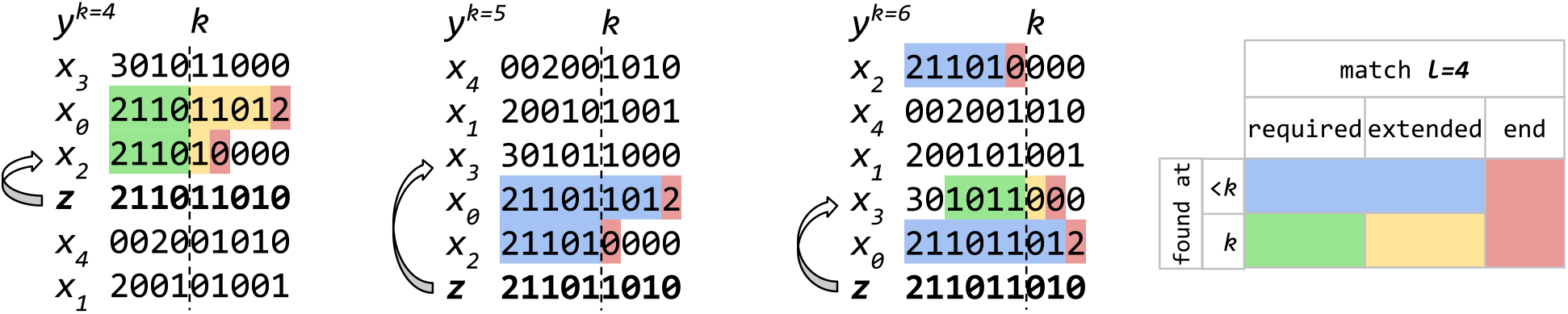
The process of finding long matches. The states of the algorithm at *k* = 4, 5, 6 are shown. The arrows indicate setting the pointer *p* to skip over the previously found matches. At *k* = 4, no matches have been previously found, so *p* is set to the sequence immediately above *z*, and the block of two matches above *z* are found. At *k* = 5, the two previously found matches are skipped, but no new matches are found. At *k* = 6, one of the previously found matches has terminated, so we skip over the remaining ongoing match to find the new match.

In genetic sequence data, recombination events result in match lengths of non-uniform distribution. To take advantage of the disproportionately large number of relatively short matches, we formulate the following heuristic: We begin by linear searching for the match end using the syllabic panel for several iterations (e.g. 10, covering 10*B* sites). If our match is among the few exceptionally long ones, we then switch to binary search with hashes for the remaining syllables. This way, we are able to find the match end in a small constant time without hashing in the average case, while bounding the runtime by *O*(log *n*) in the worst case.

#### 2.4.4 Allowing for queries in panels with unevenly distributed sites

When querying with genetic distance (cM) or physical distance (bps), site locations are nondecreasing but not necessarily uniformly distributed. With proper bookkeeping and traversal, we can query with unevenly distributed sites without affecting the time or space complexity (see Appendix 6.8). Appendix 6.2 contains the pseudocode for query search with variably distributed sites, of which site-length query search is a subproblem.

## 3 Results

We benchmarked Syllable-Query using *B* = 64 and *B* = 128 for reasons discussed in Appendix 6.3. We refer to Algorithm 3 of Sanaullah et al., the most time- and space-efficient previously published solution to the *L*-long match query problem, as the full memory algorithm. Between the static and dynamic versions of the algorithms presented with d-PBWT, we chose to implement the static version of the full memory algorithm for consistency with the static nature of Syllable-Query and because the static version is more competitive in terms of memory.

We observed the full memory usage on chromosome 21 from UK Biobank (974,818 haplotypes and 9,793 sites) to be 150.4 GB. Given that the asymptotic memory usage of the full memory algorithm is proportional to the panel size *MN*, we extrapolated the full memory requirement for querying on the 22 autosomal chromosomes from UK Biobank consisting of 974,818 haplotypes and 658,720 markers to be 10.1 TB. In comparison, Syllable-Query used only 162 GB and 91.4 GB with *B* = 64 and *B* = 128, respectively, for the same task, yielding respective memory reduction factors of 62 and 110. Figure 3 provides the memory usage reductions for every chromosome based on its size. The positive trend between memory reduction and the number of sites is due to the positive trend between the number of sites and marker density per genetic distance, allowing the syllable dictionaries *r* to use less space per syllable. Over the 22 autosomal chromosomes collectively, we observed *ρ* ≈ 28.3 for *B* = 64 and *ρ* ≈ 7.2 for *B* = 128, demonstrating that *ρ*, which is inversely proportional to the space taken by our dictionaries *r*, is likely to be of considerable magnitude for genetic sequence data due to linkage disequilibrium.

**Figure 3:**
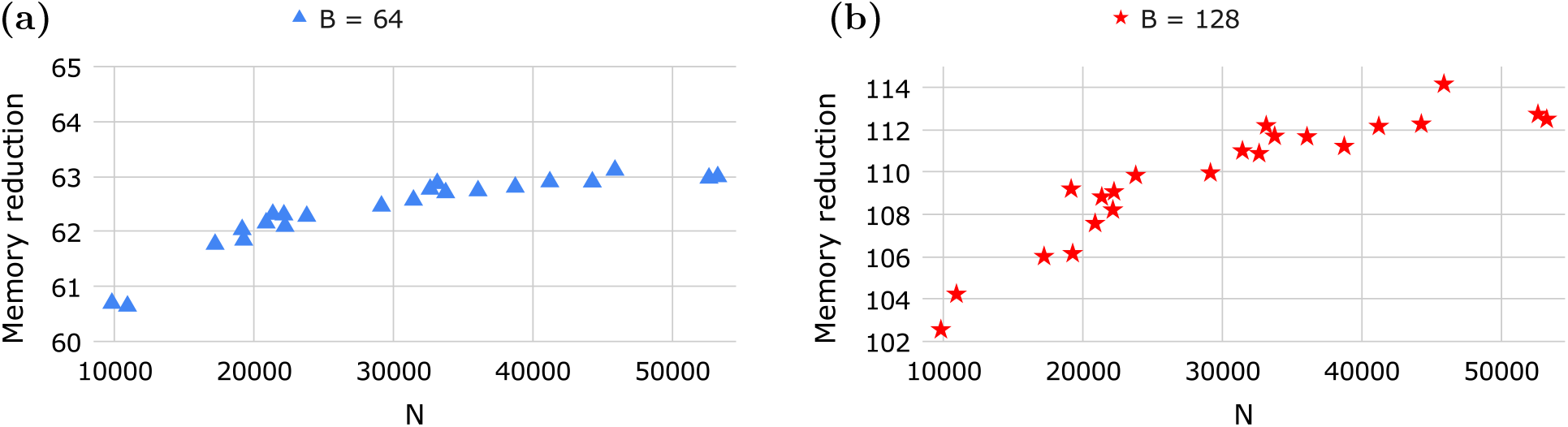
Memory reduction factors of Syllable-Query compared to the full memory algorithm versus the number of sites *N* over the UK Biobank autosomal chromosome genotype data (*M* = 974, 818). Each dot is a chromosome.

We benchmarked the runtime of Syllable-Query with respect to the number of matches, sites, and haplotypes on data from UK Biobank and the 1000 Genomes Project. In every panel, we removed the first 100 haplotypes to use as query haplotypes and recorded the average CPU runtime (on a single core of a 2.10 GHz Intel Xeon Silver 4116 Processor) and the average number of matches over the 100 queries. Our intention behind running many queries in succession was to stabilize the degree of runtime volatility due to factors such as the CPU cache, as well as to simulate the practical setting of matching a query panel against a predefined panel.

Figure 4a shows the runtime of Syllable-Query to scale about linearly with the number of matches *c* and puts it into perspective with the full memory algorithm runtime. The most observable increases in the Syllable-Query runtime trend occur when *L* drops below *kB* – 1 for some small integer *k* (see Appendix 6.9 for why). Figure 4b motivates our match extension heuristic by confirming the relative shortness of the average match length. Moreover, the comparable slopes of the trendlines in Figure 4a demonstrate that our virtual insertion and match extension heuristics are satisfactorily fast for real data (recall that the full memory algorithm is known to scale very well with the number of matches as it only processes matches upon termination). The *y*-intercepts of the trendlines further reveal that Syllable-Query is significantly faster than the full memory algorithm, even in the computationally unfavorable situations with small *L* mentioned above. We attribute the speedup in performance primarily to two reasons: (1) With *B* times fewer syllables than sites, the syllabic panel is much faster to iterate through. Although the reduced number of syllables is accompanied by a slight runtime factor *β* mentioned in Appendix 6.3, *β* only appears in the match refinement stage and therefore minimally affects runtime, evident in the only slightly higher slope of the *B* = 128 trendline than that of *B* = 64 in Figure 4a. (2) The CPU cache grants the CPU fast access to frequently used memory locations. Therefore, with less memory required by Syllable-Query, the CPU cache loads in data pertaining to more sites at a time, thereby saving time on data retrieval.

**Figure 4:**
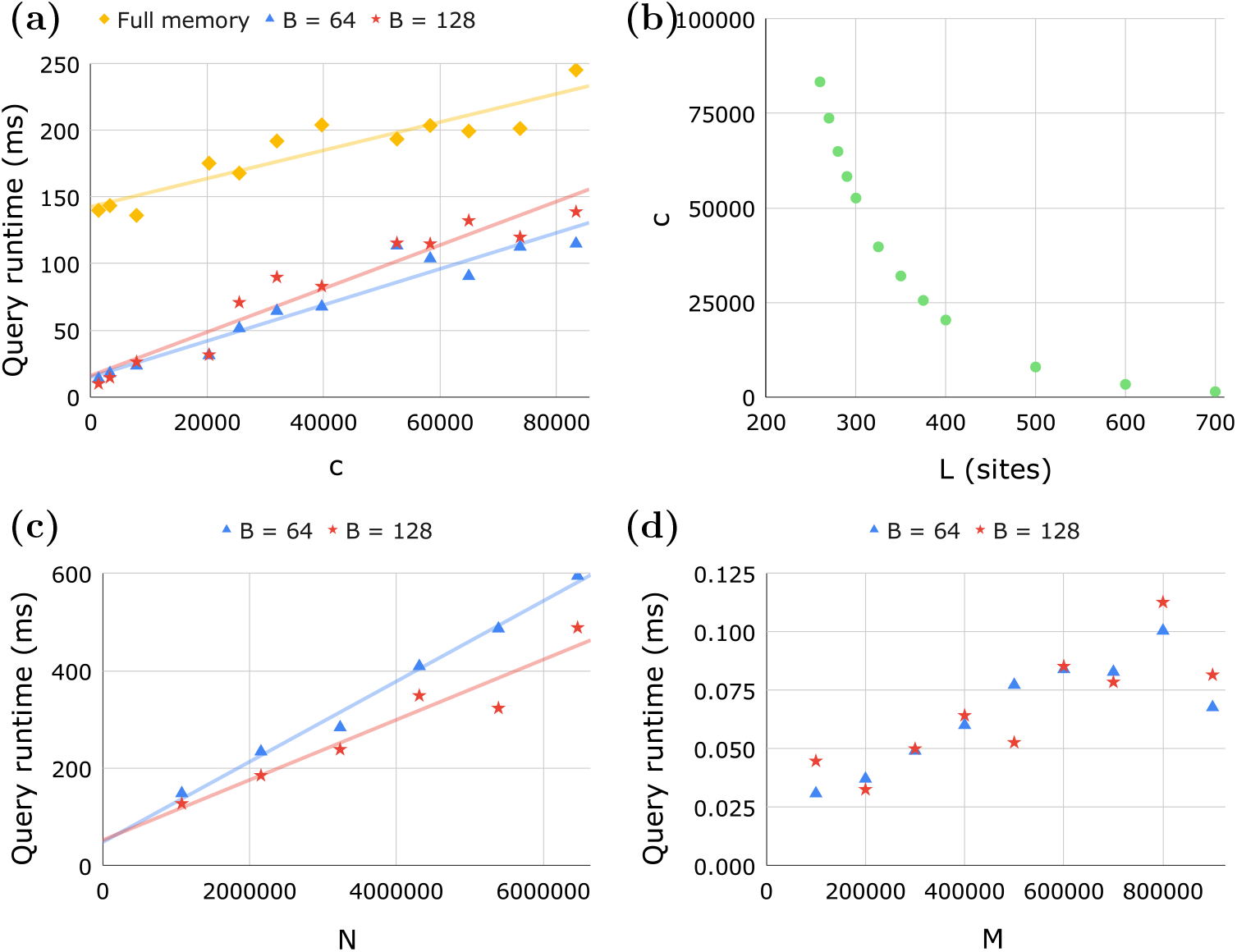
Benchmarking on UK Biobank and 1000 Genomes data. **(a)** The average query runtime versus the average number of matches *c* for each algorithm on chromosome 16 of UK Biobank (*M* = 974, 718; *N* = 23, 774). **(b)** The average number of matches *c* versus the minimum match length *L*, where the *x*-axis of part (a) corresponds to the *y*-axis of part (b). **(c)** Average query runtime versus *N* on chromosome 1 sequence data from the 1000 Genomes Project (*M* = 4, 908). The average number of matches *c* was kept at a relatively constant high (94, 000 < *c* < 102, 000) to examine the impact of the log *n* components in the time complexity. *N* was varied by choosing roughly uniformly distributed subsets of sites. **(d)** Average query runtime versus *M* on chromosome 1 of UK Biobank (*N* = 53, 260). The average number of matches *c* was kept at a relatively constant low (0 < *c* < 37) to limit variation within the time complexity term independent of *M* and to measure performance on queries with high length requirements.

For the runtime benchmarks versus *N* and *M*, we exclude comparison with the full memory algorithm due to the sheer memory consumption that would be required. Regardless, the difference in *y*-intercepts in Figure 4a suggests that Syllable-Query scales better with *N* than the full memory algorithm. Figure 4c confirms the roughly linear runtime of Syllable-Query with respect to *N*. The memory usage for the benchmark on sequence data depicted in Figure 4c (*M* = 4, 908; *N* = 6, 468, 094) using *B* = 64 and *B* = 128 was, respectively, 7.9 GB (*ρ* ≈ 61) and 4.1 GB (*ρ* ≈ 28), as opposed to an extrapolated 500 GB required by the full memory algorithm. The significantly higher *ρ* values compared to those for the UK Biobank autosomal genotype data are to be expected, since genotype data, unlike sequence data, are limited to genetic variants, giving rise to more distinct syllable values. Figure 4d shows a diminishingly positive correlation between query runtime and *M*, as to be expected by the log *M* factor involved in virtual insertion; note too that another cause of the positive correlation is the more effective CPU cache usage for smaller *M*. Due to the term log *M*, as *M* approaches infinity, our time complexity (but not the full memory algorithm’s) approaches infinity, but our benchmarking reveals this growth rate to be negligible in practice.

To verify the empirical correctness of Syllable-Query beyond the prior theoretical discussion, especially concerning the reliability of our hash function, we ran 1,000 distinct queries each for *L* in terms of sites and cM on UK Biobank data, totaling over one million matches. For every query, Syllable-Query reported the same matches as the full memory baseline algorithm.

For genealogical search, query lengths of at least 5 or 7 cM and 700 SNPs are typically chosen, as established by simulations conducted by the DTC company 23andMe [13, 14]. Despite the minimum query length required by Syllable-Query (see Appendix 6.3), we found that our site and cM requirements using deCODE genetic maps were well below these cutoffs (see Appendix 6.10), so our requirements would not limit the application of Syllable-Query to genealogical search.

## 4 Discussion and conclusions

We have presented the Syllable-PBWT framework as a space-efficient alternative to the conventional binary PBWT. The main methodological contribution of this work is the redesign of the PBWT long match query algorithm by stripping away the most memory-intensive PBWT data structures. Transforming the binary panel into a syllabic panel can be viewed as abstracting away fine detail to lighten the memory load while retaining the information required for finding long matches. Most importantly, we introduce hash arrays to underpin Syllable-Query’s ability to query without the full-panel-sized arrays *d*, *u*, *v*. Although using *d*, *u*, *v* in a transient fashion for all-vs-all matching is appropriate, making them persistent for one-vs-all query is overly memory-costly. With the hash arrays *h*, we maintain the constant runtime exhibited by d for checking whether a match is long enough. Moreover, *h* can substitute *u* and *v* for virtual insertion but with the incurrence of a small worst-case *O*(log *M* log *n*) runtime factor for binary search.

While in this paper we aimed to present the most space-efficient solution by putting all the above design elements together, it is worth reviewing their individual contributions. The biggest memory reduction comes from creating the syllabic panel and replacing the full-panel-sized *a*, *d*, *u*, *v* with syllabic versions of *a* and *h*. To reduce the size of the panel itself, we use coordinate compression to bring the overall memory reduction factor to about *B*. In addition, when memory efficiency is not the sole priority, one may mix-and-match the design elements to create simpler alternative algorithms with lower degrees of space efficiency. Appendix 6.11 further discusses alternative Syllable-PBWT data structures.

Although the primary goal of this work is to reduce the memory footprint of the long match query algorithms, some elements of our algorithms can be used for other purposes. For example, the coordinate compression in Syllable-PBWT can be a solution for lossless compression of the PBWT panel. Unlike run-length compression of the divergence array mentioned by Durbin [4] which is not friendly for real-time querying, our Syllable-PBWT data structures support regular PBWT algorithms within the compressed format without decompression. Therefore, our algorithm can also be applied to all-vs-all matching; a naive method is to query each haplotype against the rest, although there may be a more efficient method.

Conceptually, Syllable-PBWT is reminiscent of multiallelic-PBWT (mPBWT) [22] in that the sequence values are elements of a variably-sized alphabet. However, Naseri et al. [22] only presented algorithms for panel construction and all-vs-all matching in multiallelic data but none for one-vs-all query. A contribution of this work is the long match query algorithm absent from mPBWT.

Despite the utility of *L*-long matches, one drawback is their requirement for match exactness, whereas real data often contain genotyping and phasing errors. Encouragingly, the contributions in this work could be adapted to a mismatch-tolerant variation of the *L*-long match query problem. Since past efficient solutions only consider matches upon termination, little potential remains for looking past match interruptions. In Syllable-Query, on the other hand, matches are considered as soon as they reach the threshold length and are then manually extended. Therefore, the extension process can be modified to continue as long as the number of mismatches remains below a specified parameter. The various starts and ends of the fragmented match could then be recorded in an event schedule, a more intricate development of the match end tracker in our current algorithm, to swiftly bypass the fragments composing previously found matches.

Beyond methodological contributions, we showed that Syllable-PBWT and Syllable-Query delivered a memory reduction factor of over 100 in real sequences from the 1000 Genomes Project and the UK Biobank. For UK Biobank, while the state-of-the-art query algorithm [15] requires 10 TB of memory, Syllable-Query only requires 91 GB. This innovation will allow online genealogical search to be conducted with much more modest hardware and on even larger data sets in the future.

## 5 Acknowledgements

This work was supported by the National Institutes of Health grants R01 HG010086 and R56 HG011509. This research has been conducted using the UK Biobank Resource under Application Number 24247.

## 6 Appendix

### 6.1 Syllable-PBWT pseudocode

Algorithm 1 describes the construction of the Syllable-PBWT data structures.

#### Algorithm 1: Construction of Syllable-PBWT data structures

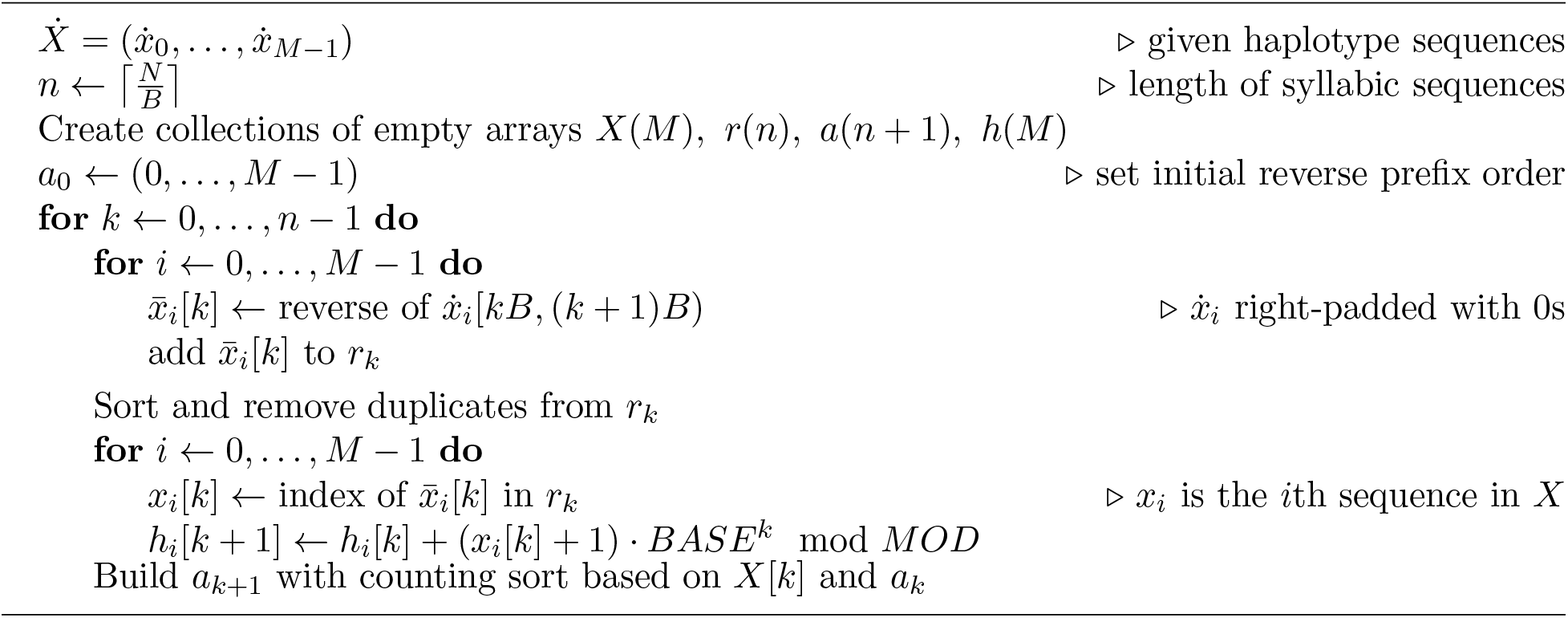

### 6.2 Syllable-Query pseudocode

Algorithm 2 describes the Syllable-Query procedure for query search with variably distributed sites. To query in units of sites, we would use a simplified version of Algorithm 2 in which the *q* pointer is always *k* – *l*, where 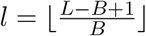.

### 6.3 Choice of *B*

Several considerations must be made for choosing an appropriate value for *B*.

- The value of *B* restricts the smallest allowable query length because if a small query length *L* causes *l* = 0, then the runtime would skyrocket. If we were to override *l* to be 1, then matches that do not extend over at least one full syllable would pass unseen by our algorithm. Specifically, in the worst case scenario, a match may nearly entirely span two adjacent syllables (2*B* – 2 sites), but we would be unable to detect it. If, however, the match is any longer, it must span at least one full syllable, meaning that 2*B* – 1 sites is the smallest query length that a given *B* can handle. Letting *B* be 64 or 128 leads the smallest query length to be 127 or 255 sites, respectively, which are well below the query lengths used to detect IBD segments in practice. Similarly, for variably distributed sites (i.e. when querying with genetic or physical distance), the query length must be greater than the longest distance spanning exactly two adjacent syllables. See Appendix 6.10 for evidence of these *B* values yielding genetic query length requirements that comfortably allow for IBD segment detection. Further increasing *B* would be accompanied by the risk of limiting practical use.
- In order to store and operate on the raw syllable values, the programming language we use for implementation must provide data types to support a *B*-bit integer. The GNU C++ compiler provides the `unsigned long long` data type which supports *B* = 64, and on 64-bit processors, it also provides the `unsigned __int128` data type which supports *B* = 128. These built-in data types motivated our use of these *B* values, although in theory, the Boost C++ libraries could support even higher values of *B*.
- The predominance of 64-bit processors in recent decades has allowed for typical instructions involving 64-bit data to be executed within a single CPU clock cycle [23, 24]. Choosing *B* = 64 would allow us to make most efficient use of this architecture, while choosing *B* = 128 would introduce a small constant runtime factor to bitwise operations between raw syllable values. In general, the runtime factor of basic operations on raw syllable values for a chosen *B* is 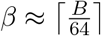, because while instructions involving *B*-bit data (*B* > 64) do not necessarily require multiple clock cycles, such is the worst case scenario.
- The final motivation for not further increasing *B*, even though doing so would lead to more memory reduction, is the diminishing rate at which increasing *B* increases the memory reduction factor. The rate is diminishing because as *B* increases, raw syllable values are more likely to be distinct, so the ratio *ρ* will decrease, causing the memory usage of *r* to decrease evermore slowly as *B* increases.

#### Algorithm 2: Syllable-Query to report *L*-long matches between query haplotype and panel

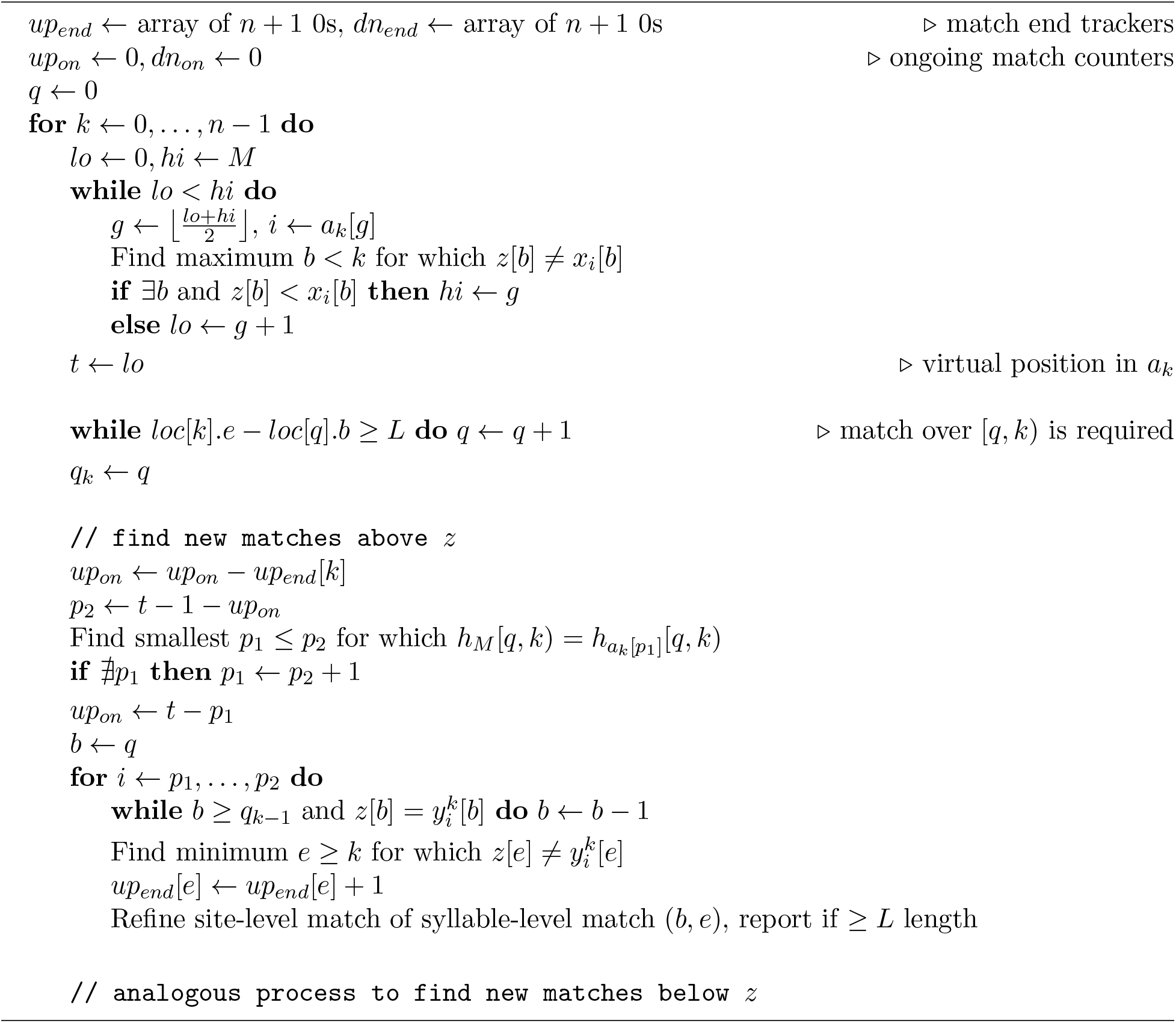

With the factors discussed above in mind, we implement Syllable-Query with *B* ∈ {64, 128}.

### 6.4 Reliability of the polynomial hash function

To create a strong hash function with minimal chance of collisions, several conditions must be satisfied: the modulus should be a large prime, the polynomial variable should be greater than the largest value in the input sequence, and the modulus and polynomial variable should be relatively prime [21, 18]. In our implementation, we set *BASE* = 2^48^ – 59 and *MOD* = 2^64^ – 59, satisfying these criteria (as well as bounding every hash to fit within a 64-bit integer), so our polynomial hash function can be considered a pseudorandom function through which two identical inputs are guaranteed to yield the same hash and two distinct inputs have a 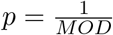 chance to yield the same hash. To illustrate the theoretical robustness of the hash function, suppose that we were to perform *q* = 10^10^ independent comparisons of distinct sequences. The chance *P*(*p*, *q*) = 1 – (1 – *p*)^*q*^ of a positive number of collisions occurring and the expected number *E*(*p*, *q*) = *pq* of collisions would both be less than 10^−9^, demonstrating that even with a number of distinct-sequence comparisons that dwarfs what would be required in a practical setting, the expected chance of a false positive remains negligible.

### 6.5 Refining match boundaries

To obtain the single-site resolution of the match boundaries in *O*(*β*) time, we perform bitwise operations on the raw syllable values where a match (non-inclusively) starts and ends. The GNU C++ compiler provides the bitwise XOR operator ^^^ (which gives an integer in which every bit is 0 for which the two operands have the same value) and the functions `__builtin_clzll` and `__builtin_ctzll` which count the number of leading and trailing 0s, respectively, in a variable of the 64-bit data type `unsigned long long`. These functions can be used to count the number of leading or trailing 0s in the bitwise XOR between two integers, giving the length of the suffix or prefix match, respectively, of the reverse binary representations of the integers (recall that the raw syllable values are reversed binary haplotype substrings parsed as binary numbers). For *B* = 128, we see *β* come into play because the left and right 64-bit halves of the 128-bit integer have to be operated on separately, so the runtime of refining the 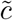 potential long matches is 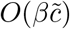.

### 6.6 Motivation behind searching for ongoing long matches

A convenient characteristic of the long match query problem for binary sequences is that long matches that terminate at a given site *k* all occur in a single contiguous block in *a*_*k*+1_, because for any allele value *ż*[*k*], there is exactly one allele value not equal to *ż*[*k*]. Past solutions have made use of this convenience by maintaining pointers for the block in *a_k_* of long matches to efficiently obtain the block in *a*_*k*+1_ of newly terminated matches, thereby counting every long match once and only once. In long match query with Syllable-PBWT, on the other hand, there could be up to *M* – 1 blocks in which terminated matches lie, implying the need to consider matches before termination.

### 6.7

#### Proof of Lemma 1

*Proof*. In Algorithm 2 of PBWT, Durbin observes that the start *b* of the longest ongoing match at site *k* between 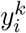 and 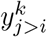 is max_*i*<*m*≤*j*_*d_k_*[*m*], where the divergence array *d_k_* is defined such that *d_k_*[0] = *k*, and for *i* > 0, *d_k_*[*i*] is the smallest *b* for which 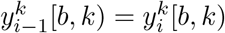. With this observation, 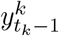 must be the sequence with which (1) *z* has the longest ongoing match, (2) among sequences with a reverse prefix over syllables [0, *k*) that is lexicographically less than that of *z*. Condition (1) must be satisfied because no other match above *z* can spontaneously obtain an earlier start, which by definition is not earlier than that of the match with sequence *x_i_*. Condition (2) must be satisfied because the match persisting as *k* increases implies that the same values are being prepended to the reverse prefixes of *z* and *x_i_*, thereby not affecting their lexicographical ordering. Even if sequence *x_i_* has an equally long match with *z* as another sequence *x_j_* above *z*, the construction of the positional prefix array based off that of the previous syllable guarantees the retention of the relative positions of sequences between which a match persists. Therefore, until the match ends, both conditions will remain satisfied, implying that *a_m_*[*t_m_* – 1] = *i* at every syllable *k* ≤ *m* ≤ *k* – *l* + *s*.

### 6.8 Allowing for queries in panels with unevenly distributed sites

While for the site-length query, potential matches had to match over syllables [*k* – *l*, *k*), we must now maintain an index *q* that is incremented at every syllable *k* until any potential match must match over syllables [*q_k_*, *k*), where *q_k_* denotes the value of *q* at syllable *k*. Let *loc*[*k*].*b* and *loc*[*k*].*e* be the locations of the first and last sites, respectively, of syllable *k*, and generalize *L* to be the query length in the same units as in *loc*. At every syllable *k*, we update *q* by incrementing it until *loc*[*k*].*e* – *loc*[*q*].*b* < *L* so that no partial match at syllable *k* will make up for *q* being an invalid match start; a full match at syllable *k* implies that the match will be reconsidered at syllable *k* + 1. Note that it is guaranteed that *q_k_* < *k* (and therefore potential matches must span at least one full syllable) if the query length is greater than the maximum distance spanning exactly two adjacent syllables (see Appendix 6.3 for details).

If we were dealing with single-site resolution data, we would be done, but instead our *B*-site resolution *q* index does not suffice for finding the match start. For example, suppose that *loc*[*k*].*e* – *loc*[*k* – 1].*e* is significantly greater than the corresponding distance between any two preceding adjacent syllables. This would lead the gap *q_k_* – *q*_*k*_1_ to be large, leaving the match starts between syllables *q*_*k*_1_ and *q_k_* untracked. To backtrack efficiently, first assume that using the same approach as for the site-length query, we have determined the block of new matches above *z*, e.g. from 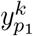 to 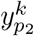 where *p*_1_ < *p*_2_, spanning syllables [*q_k_*, *k*). As we now iterate *i* from *p*_1_ to *p*_2_, we can decrement a running pointer *b* starting from *q_k_* to find every match start. According to Lemma 2, maintaining *b* would not affect the query time complexity.

#### Lemma 2.

*The total number of times the pointer b must be decremented for a given query is bounded by n*.

*Proof*. According to Durbin’s observation stated in Appendix 6.7, the reverse prefix match with *z* over syllables [0, *k*) must be at least as long for 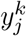 as for 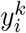 if *i* < *j* < *t_k_*. Therefore, at a given syllable *k*, the non-inclusive match start pointer *b* can be updated for a given 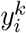 by simply decrementing *b* until 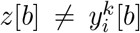. Furthermore, the fact that these new matches are found at syllable *k* rather than earlier means that their match starts must be greater than *req*_*k*_1_, so *b* may be decremented at most from *req_k_* to *req*_*k*_1_ at syllable *k*. Thus, the total number of decrementations necessary to backtrack to the match starts for a given query is bounded by the telescoping sum ∑_0<*k*≥*n*_ *req_k_* – *req*_*k*–1_, which is at most *n*.

### 6.9 Syllable-Query runtime on small *L*

To see why the most observable increases in the Syllable-Query runtime occur when *L* drops below *kB* – 1 for some small integer *k*, recall that for Syllable-Query to identify all matches spanning *L* sites, it must consider all 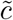 potential long matches spanning 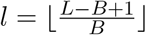 syllables, which means that, for example, when *L* < 3*B* – 1, matches spanning a mere syllable must be considered, whereas when *L* = 3*B* – 1, only matches spanning at least 2 syllables are potential matches. These runtime spikes are only expected to be considerable for relatively small *L*, since genetic recombination leads the absolute rate of change of *c* with respect to *L* to start large before drastically dropping off, as shown in Figure 4b.

### 6.10 Syllable-Query length requirements

To confirm the practicality of Syllable-Query despite the restriction on query length (see Appendix 6.3 for details), we measured the smallest queryable genetic length (SQGL) in cM for each autosomal chromosome with the deCODE genetic map [25] in UK Biobank data. For *B* = 64 and *B* = 128, the smallest queryable site lengths are 127 and 255, respectively. The highest SQGL for 127 and 255 sites was 2.7 and 4.2 cM, respectively. The average SQGL for 127 and 255 sites was 2.2 and 3.3 cM, respectively. These requirements are well below the query lengths of 5 or 7 cM and 700 SNPs utilized by 23andMe for genealogical search [13, 14]. Since UK Biobank has a relatively low marker density, our SQGLs will likely also support genealogical search on biobanks in general. Unless especially small *L* values are to be used, we recommend choosing *B* = 128 for greater memory reduction, given the similar runtimes between *B* = 64 and *B* = 128 shown in Figure 4.

### 6.11 Alternative Syllable-PBWT data structures

#### 6.11.1 Inverse positional prefix array

In the presented version of Syllable-Query, two nested binary searches with hashes were necessary to find the virtual position *t_k_* of *z*. By storing the inverse 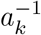 of every positional prefix array *a_k_*, a single binary search over the pairs (*x_i_*[*k*], 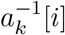) can find *t_k_*, where 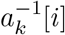 can be thought of as the position of haplotype *x_i_* in *a_k_*.

#### 6.11.2 Divergence array

Although the most efficient version of Syllable-Query replaces *d* with *h*, other applications of Syllable-PBWT may wish to utilize the divergence arrays *d*. As such, we present the construction for the divergence arrays in a syllabic panel. The divergence array *d_k_* is built alongside *a_k_* and stores, for every pair of adjacent sequences in *a_k_*, the starting index of their longest reverse prefix match. Formally, *d_k_*[0] = *k*, and for *i* > 0, *d_k_*[*i*] is the smallest *b* such that 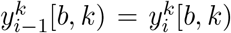. In Algorithm 2 of PBWT, Durbin observes that the start *b* of the longest ongoing match at site *k* between 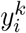 and 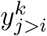 is max_*i*<*m*≤*j*_ *d_k_*[*m*]. Using this observation and the binary nature of allele values, Durbin is able to compute *d*_*k*+1_ in *O*(*M*) time, given *a_k_*, *d_k_*, and *X*[*k*], by maintaining two running maximums of *d_k_* values.

We seek to compute *d*_*k*+1_ for sequences with up to *M* possible syllable values. When *i* = 0 or 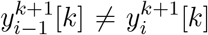, we simply have *d*_*k*+1_[*i*] = *k* + 1. Otherwise, let us define 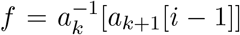 and 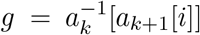. In other words, given that *i* – 1 and *i* are positions in *a*_*k*+1_, *f* and *g* are their respective positions in *a_k_*. Considering Durbin’s observation, we then have *d*_*k*+1_[*i*] = max_*f*<*m*≤*g*_*d_k_*[*m*]. To efficiently perform the range maximum query, we utilize the sparse table data structure. At every syllable *k*, we build the sparse table off of *d_k_* so that *d*_*k*+1_ can be built off of the sparse table. By splitting *d_k_* into blocks of size ⌊log *M*⌋, the sparse table can be built on the maximums within the blocks in *O*(*M*) time and space. After construction, the *M* potential queries each take *O*(1) time if *M* < 2^64^ and *O*(log *M*) time otherwise. Since the sparse table is replaced at every syllable and discarded after precomputation, it does not contribute to the overall memory usage. Therefore, for *M* < 2^64^, *d* requires *O*(*Mn*) time to compute and *O*(*Mn*) memory to store.

